# K-MARVEL: K-Mer based Antimicrobial Resistance Virtual Exploration Lab

**DOI:** 10.1101/2025.10.08.681127

**Authors:** Nirmal Singh Mahar, Sverre Branders, Manfred G. Grabherr, Ishaan Gupta, Rafi Ahmad

**Author notes:** Corresponding Author, Correspondence of Ishaan Gupta, Correspondence of Rafi Ahmad.

## Abstract

The rapid global spread of antimicrobial resistance (AMR) necessitates a new generation of computational tools for its surveillance. While next-generation sequencing offers unprecedented insight into the resistome, current methods face a trade-off: assembly-based approaches are computationally expensive and struggle with complex metagenomes, whereas direct-mapping of long reads is hampered by high error rates that obscure critical resistance-conferring mutations. Here, we present K-MARVEL (K-Mer based Antimicrobial Resistance Virtual Exploration Lab), a novel, open-source method to capture ARGs and resistance-conferring mutations from short and long-read sequencing datasets. It operates in protein k-mer space, providing inherent tolerance to nucleotide-level sequencing errors. On a comprehensive benchmark of 61 long and 49 short-read diverse datasets, K-MARVEL demonstrated superior accuracy, achieving F1-scores of 0.9783 and 0.9754 for short and long-read datasets, respectively. Its implementation in Rust enables high speed through parallelization while guaranteeing memory safety. Computationally, it demonstrated superior performance to conventional assembly-based methods, achieving an average speed up of 7x on short-read datasets and 5x on long-read datasets. In terms of memory footprint, it outperformed the assembly-based approaches for short-read datasets, but its memory footprint was comparable for long-read datasets. Notably, K-MARVEL accurately reconstructs functional genes from genomically fragmented evidence, providing a more comprehensive resistome assessment. In conclusion, K-MARVEL provides a scalable, flexible and memory-efficient solution for AMR surveillance. Its unique capabilities for handling noisy long-read data and complex genomic scenarios make it a powerful tool for researchers and public health scientists.

K-MARVEL is open-source and freely available at https://bitbucket.org/amr-avenger/k-marvel under the GPL version 3 license.

## Introduction

Antimicrobial resistance (AMR) represents a critical and escalating global health threat, diminishing the efficacy of conventional antibiotics and complicating the treatment of bacterial infections^1^. The proliferation of AMR is primarily driven by the misuse and overuse of antimicrobial agents, which selects for and facilitates the dissemination of antibiotic resistance genes (ARGs) within microbial populations^2^. Consequently, the precise identification and characterization of ARGs are fundamental to elucidating resistance mechanisms, monitoring their dissemination, and informing the development of novel therapeutic strategies^3,4^.

The advent of high-throughput sequencing technologies has revolutionized AMR surveillance, enabling the comprehensive detection of ARGs directly from complex metagenomic and clinical samples, unconstrained by the limitations of the conventional targeted molecular assays^5,6^. Standard bioinformatic methods for identifying ARGs in sequencing data typically involve direct read alignment to curated ARG databases or a more computationally intensive assembly-first approach, where reads are assembled into contigs before database mapping^6^. While the latter method mitigates challenges associated with short read lengths, its significant computational burden and time requirements often hinder rapid analysis.

Third-generation sequencing technologies, such as Oxford Nanopore Technologies (ONT), address the read-length limitation by generating long reads that can span complete genes in real-time, thereby simplifying read-based mapping^7,8^. However, the high error rate inherent to these technologies often obscures the detection of specific point mutations, which is crucial for identifying point mutations responsible for resistant chromosomal genes (e.g., in *gyrA, parC, rpoB*)^9^. Hybrid error-correction strategies, which leverage accurate short reads to polish error-prone long reads, or *de novo* assembly, can improve accuracy. Nevertheless, these solutions are suboptimal for real-time applications due to the associated costs (hybrid sequencing), high computational complexity, and the requirement for sufficient sequencing depth^10,11^. Furthermore, in culture-free metagenomic samples, assembly is complicated by host DNA contamination, heterogeneous species abundance, and diverse microbial backgrounds^12,13^. Also, one major challenge in AMR surveillance is that current computational methods cannot simultaneously detect acquired resistance genes and resistance-conferring mutations without significant limitations. Methods like Resistance Gene Identifier (RGI) depend on prior genome assembly^14^, while tools like PointFinder require the user to specify the bacterial strain in advance, hindering the analysis of complex or unknown samples^15^.

To overcome these limitations, we present a novel, alignment-free method for directly detecting ARGs and point mutations from raw sequencing reads. We introduce K-MARVEL (K-Mer based Antimicrobial Resistance Virtual Exploration Lab), an open-source software for efficient and accurate ARG profiling from short- and long-read whole genome or metagenome sequencing data. K-MARVEL employs a specialized k-mer matching and extension strategy in protein space, bypassing the need for prior assembly or taxonomic classification. This allows for the simultaneous detection of acquired ARGs and resistance-conferring mutations in chromosomal genes.

We demonstrate that K-MARVEL outperforms assembly-dependent methods in computational efficiency and provides highly accurate results across multiple publicly available datasets, encompassing long and short-read technologies at varying sequencing depths.

## Methods

K-MARVEL is a high-performance computation tool, implemented in Rust, for identifying Antimicrobial Resistance Genes (ARGs) from DNA sequencing data. The tool is optimized for both short-read and long-read sequencing technologies. The workflow consists of four modules, each of which is described below. The schematic overview of the modules involved in K-MARVEL is presented in Figure 1.

**Figure 1:**
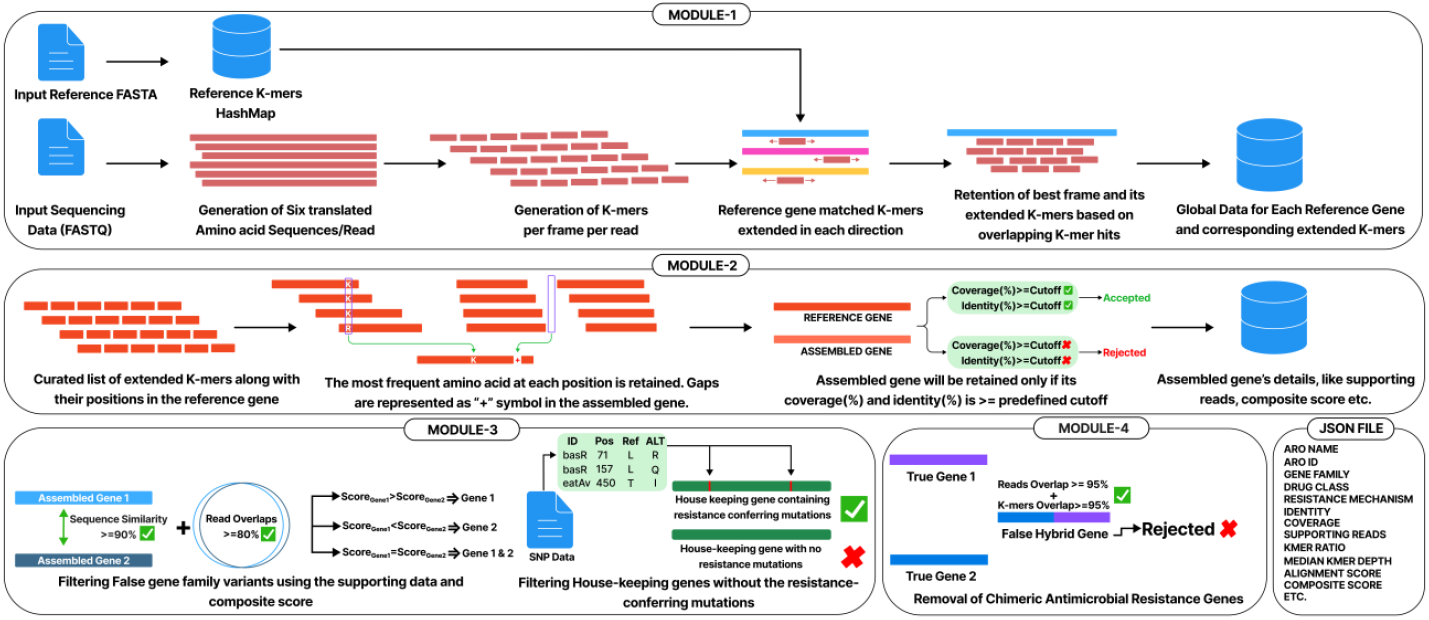
Schematic of the K-MARVEL Algorithm. K-MARVEL is divided into four modules to capture ARGs from raw sequencing data. (1) Read translation and k-mer generation, followed by best frame selection, (2) ARG assembly and filtering by coverage/identity cutoffs, (3) filtering of false gene family variants and house-keeping genes without the resistance-conferring mutations, and (4) removal of false chimeric ARGs. The final output provides detailed information about the assembled genes in JSON format.

**Figure 2.**
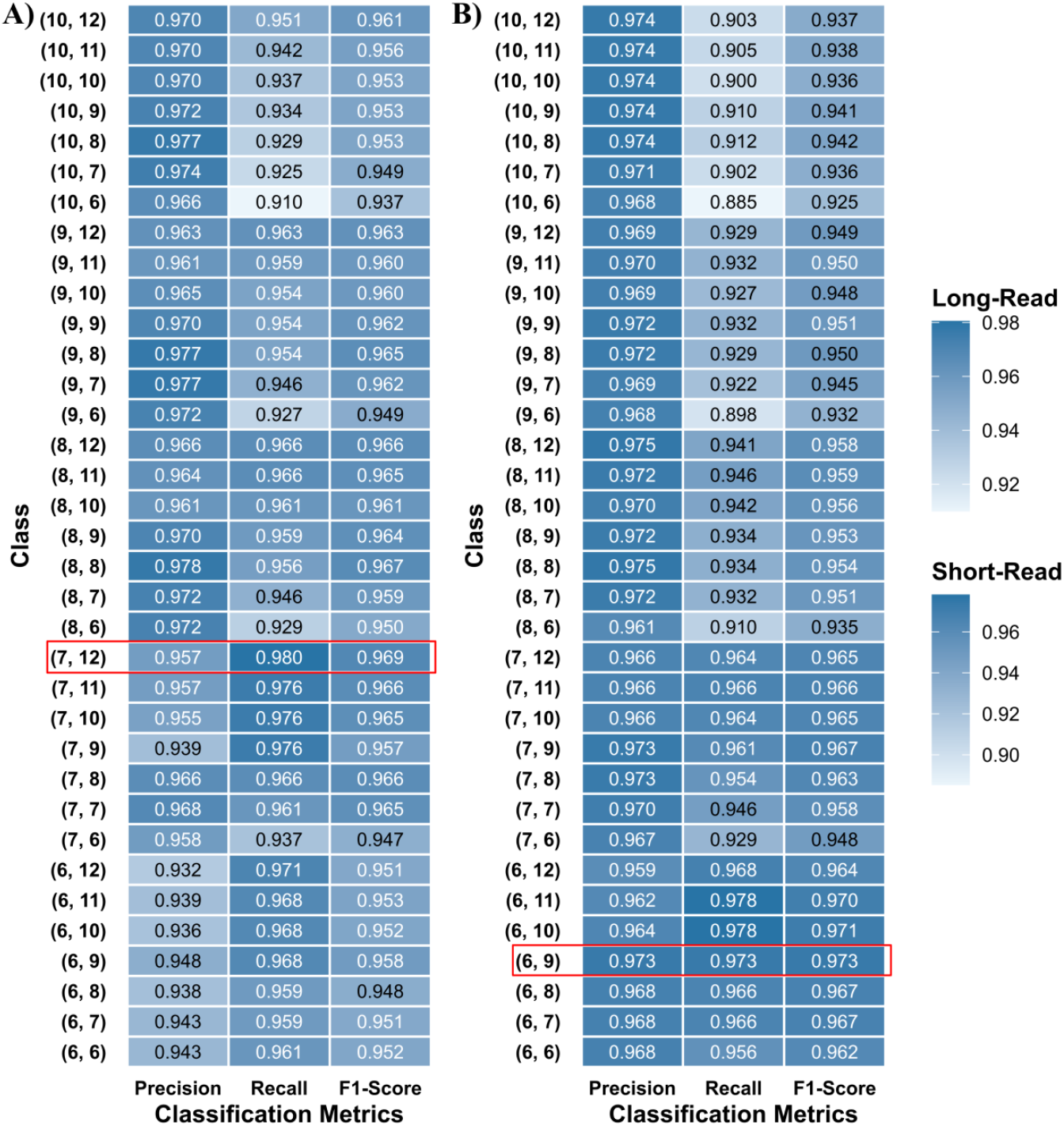
Heatmap of classification performance metrics across different combinations of kmer and extension size used for running K-MARVEL across twelve datasets. **A)** Long read sequencing datasets. **B)** Short read sequencing datasets. The highlighted block represents the best kmer and extension size combination with the highest F1 score. K-mer size of 7 and extension size of 12 was the best combination for long-read sequencing data, and 6 and 9 for short-read sequencing data

### Module 1: Reference Database Indexing and Read Mapping

The tool starts by indexing a reference database of known ARG protein sequences provided by the user. The protein sequences are indexed into a hash map with unique k-mers of size *k* as keys and sequence headers as values.

Given a reference amino acid sequence of length *n*,

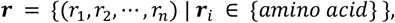

generate the set of possible k-mers of length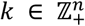, *k* ≤ *n* as

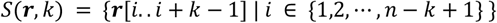

The FASTQ file provided by the user is processed in a streamlined fashion to maintain a low memory footprint. For each read, the DNA sequence is translated into all six translated frames (considering reverse complementarity). Given a DNA sequence of length *n*,

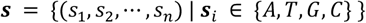

and its reverse complement

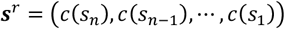

where *c*(***s***^*i*^) is the complementary nucleotide of **s**^*i*^, for each frame *f* ∈ {0, 1, 2}, naively translate each codon into its amino acid using codon table ***T***: {***A***, *T, G, C*}^3^ → {amino acid}. Given *m* valid codons in frame *f*:

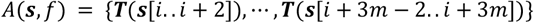

and

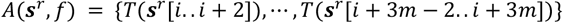

The six translated frames are converted into k-mers of size *k* as described. These k-mers are then matched with the reference database, and if any k-mers are perfectly matched with a reference k-mer, they are extended by an extension 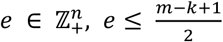 on both ends. Given a matching k-mer

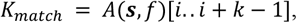

the extended k-mer

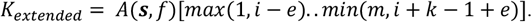

To reduce spurious k-mer matches, only those reads that provide at least five exact k-mer matches for a reference gene are kept for further analysis. Information regarding the extended k-mers, including read ID, frame ID and frame positions, is stored for further analysis.

### Module 2: Assembling ARGs

Since only one frame can encode for the gene, only the best (read id, frame) pair is kept based on the number of matchings overlapping extended k-mers. Given two extended k-mers, *K*_1_ and *K*_2_, matching a given reference gene sequence ***r*** of length *n*, their ranges are determined as the absolute positions in that sequence where they match. Let the matching range of *K*_1_ on ***r*** be denoted as

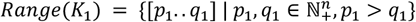

and the matching range of *K*_2_ on ***r*** be denoted as

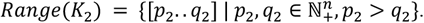

Then *K*_1_ and *K*_2_ are considered overlapping if the following conditions are met.

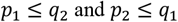

The number of overlapping extended k-mers is counted for each frame, and the frame with the highest number of overlapping k-mers is retained for the downstream analysis. If none of the frames have at least five overlapping extended k-mers, the read is excluded from further analysis of the target resistance gene.

Once all extended k-mers are curated for the reference gene, they are utilized to calculate the gene coverage. Gene coverage is computed as the percentage of amino acids in the reference gene covered at least once by the matching k-mers. Given the total number of amino acids covered at least once as *n*_*covered*_, the gene coverage is

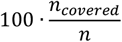

The assembly step will be done only if the gene coverage is more than or equal to the pre-defined coverage threshold.

A consensus protein sequence corresponding to the target gene is constructed using the retained extended k-mers in the assembly step. All extended k-mers are arranged based on their positions in the reference gene sequence and tally a vote for each amino acid based on the reference gene position. The amino acid with the highest occurrence counts at each position forms the consensus protein sequence. Suppose at any position, more than one amino acid has the same occurrence count, and one of the amino acids is the reference amino acid. The reference amino acid will be retained for the position in that case. Otherwise, the amino acid with the highest frequency/depth will be retained for the given position. In the rare case of equal depth, all the possible amino acids in that location will be stored and shown in the final output file. The reference gene location for which no amino acid can be curated due to the absence of reference k-mers is represented by the “+” symbol, i.e. gaps in the assembled sequence are represented by the “+” symbol. To be concord with the assembly-based ARG detection tools, the identity (%) is calculated by employing the Smith-Waterman-based Local alignment with BLOSUM62 scoring metrics provided by the rust-bio crate version 2.3.0^16^. The assembled genes with an identity (%) greater than or equal to the pre-defined cutoff set by the user will be retained for further analysis.

To filter the false variants in the downstream analysis, a composite score is derived from (I) the sequence identity to the original reference gene sequence, employing the Needleman-Wunsch algorithm provided in rust-bio crate version 2.3.0^16^; (II) kmer ratio, defined as the ratio of reference k-mers captured in the data and the total number of possible reference k-mers *n* −*k* + 1; and (III) normalized alignment score, defined as the alignment score calculated by the global alignment normalized by the reference gene length. The formula for the composite score is provided in equation 1.

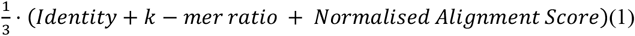

### Module 3: Filtering False Variants and House-keeping genes

Because of the high sequence similarity between the members of the same gene family, there is a possibility of assembling false variants. These variants are filtered based on the two-step strategy.

Our original assumption was that any read would encode for only one gene, but this is not true in the case of long reads, in which the same set of reads might encode the neighbouring genes. Hence, in the first step, the tool will check for the overlapping reads between the gene pair. If the overlapping reads are more than equal to 80% of the reads supporting the gene, then the tool will look for the sequence similarity between the assembled sequence of the gene pair. We employ the algorithm to calculate the longest common subsequence between two strings for sequence similarity. If the longest common subsequence is at least 90% of the smallest gene in the pair, the gene with the highest composite score will be retained. If both genes in the pair have the same composite score, then the tool will retain both.

Certain housekeeping genes/structural genes also confer antibiotic resistance only in the presence of specific mutations. To capture these mutated housekeeping genes, the tools perform a secondary screening based on the curated set of resistance-causing mutations provided by the user. Genes containing at least one curated mutation from the known set of mutations will be retained for further analysis.

### Module 4: Removal of Chimeric Antimicrobial Resistance Genes

In samples in which multiple variants of the same gene family exist, there might be a possibility of assembling a chimeric gene by the tool, which is assembled by the extended k-mers of the true variants of the gene family. We leverage the extended k-mers and supporting reads supporting the genes in a two-step process to remove these chimeric false variants. In the first step, the tool groups assembled ARGs based on their gene families and calculated the unique kmer percentage for each gene. In the case of a gene with 95% of its supporting k-mers shared with the other members of the gene family, the tool calculates the unique supporting reads percentage for the gene. If both the unique k-mer percentage and unique supporting reads percentage are less than 5%, the tool will discard the gene on the grounds of being a chimeric gene.

After running the final module, the tool will produce the resulting output as a JSON file. The main reason behind outputting the data in JSON format is to ease the tool’s integration with the other pipelines. Each gene in the JSON data will have the information regarding the assembled gene sequence, mutations found in the assembled gene, and other relevant metadata like gene family, drug class, etc.

### Calculation of optimized K-mer and Extension size for Short and Long-read sequencing data

To capture the optimum kmer and extension size for the sequencing data, we employed twelve samples, each representing a unique species for which short-read and long-read sequencing data and reference-level genome assembly were present in the public repository. To curate the ground truth set of ARGs present in these twelve reference assemblies, we ran RGI version 6.0.3^14^ with the ‘--include-loose’ parameter and ABRicate version 1.0.1^17^ with the CARD database, along with NCBI Prokaryotic Genome Annotation Pipeline (PGAP) build 7983^18^. ARGs with identity (%) >= 85 and coverage (%) >= 90 were filtered from the RGI output for further analysis. The ground truth set for a sample contained the ARGs captured by either RGI or ABRicate and supported by PGAP annotation. In the case of RGI and ABRicate capturing different variants of a gene family, the one supported by PGAP was considered in the ground truth set.

Once the samples’ ground truth set was finalized, the tool was executed with the kmer size ranging from 6 to 10 and an extension size of 6 to 12 with an identity (%) cutoff set to 85 and coverage (%) cutoff set to 90. The True Positives (TP), False Positives (FP) and False Negatives (FN) were calculated for each pair of the kmer and extension. These metrics were aggregated by the kmer and extension size for all twelve samples, and the F1 score (equation 4), the harmonic mean of Precision (equation 2) and Recall (equation 3), was calculated based on the aggregated TP, FP and FN. The kmer extension pair with the highest F1-score was selected as the sequencing technology’s default kmer and extension size.

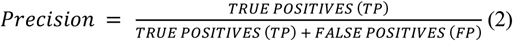

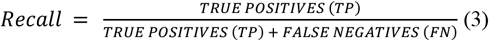

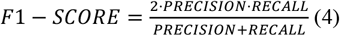

## Results

### Optimization of parameters for different sequencing technologies

To determine the optimal kmer and extension size for both the sequencing technologies, we performed a systematic grid search across the k-mer size ranging from 6 to 10 and extension size ranging from 6 to 12.

We evaluated the tool’s classification performance on twelve samples for which both short and long reads and reference-level assembly were available in the public domain. For each combination, we aggregated the classification metrics across twelve samples, and the F1-score (equation 4) was used as the primary metric for determining the optimal combination of kmer and extension size. The classification metric for each combination per sequencing technology is visualized in Figure 1. Based on the grid search, we found that the combination of 7 and 12 gave the best performance for the long-read sequencing dataset and the combination of 6 and 9 for the nshort-read data. The details about the ground truth set, captured ARGs and classification metrics per sample are provided in supplementary table 1.

### Evaluation of classification performance on empirical datasets

After the parameter optimization, we expanded the tool’s testing on other publicly available datasets. We tested K-MARVEL on 64 new datasets for long-read and 37 datasets for short-read datasets. All the datasets had reference-level genome assembly available, which was used to call the antimicrobial resistance genes (ARGs) that served as the ground truth for the comparison (Refer to methods for the detailed information about the ground truth selection). Information about the publicly available datasets utilized in this study is available in Supplementary Table 2. The classification metrics for the comparison are shown in Tables 1 and 2 for long-read and short-read datasets, respectively. K-MARVEL achieved an F1-score of 0.9783 and 0.9754 for long-read and short-read sequencing datasets, respectively, showing its universal application for short and long read sequencing technologies. Detailed information regarding the datasets used, ground truth, and observed ARGs is available in Supplementary Table 3.

**Table 1:** Global Classification performance metrics of K-MARVEL for Long-read sequencing datasets with optimized kmer and extension size (7,12).

| Number of Datasets = 61 |  |  |  |
| --- | --- | --- | --- |
| Class Designation |  | Ground Truth |  |
|  |  | Positive | Negative |
| Predicted Class | Positive | 2974 | 87 |
|  | Negative | 45 | 0 |
| Classification Metrics | Precision = 0.9716 | Recall = 0.9851 | F1-Score = 0.9783 |

**Table 2:**
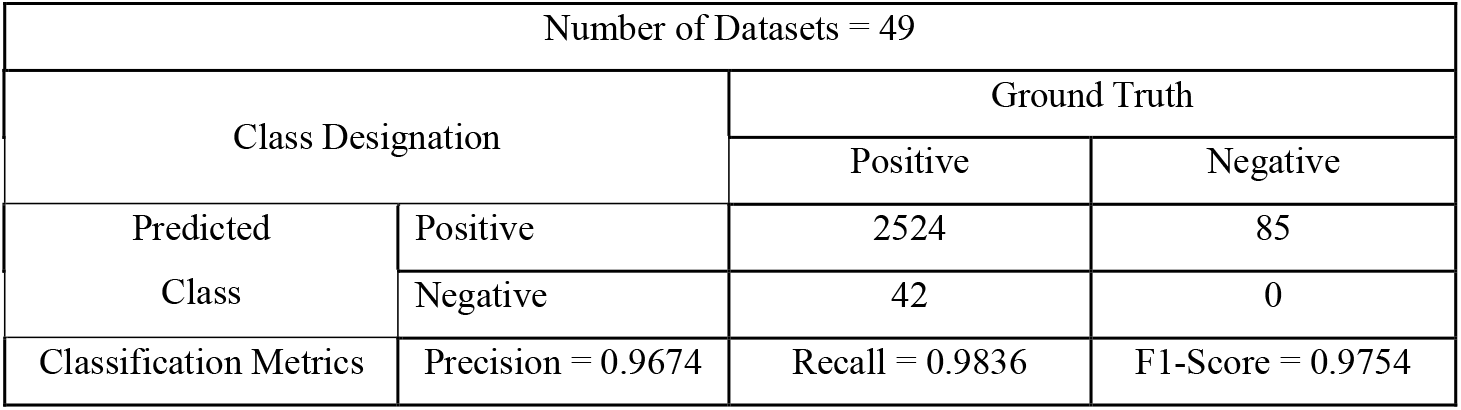
Global Classification performance metrics of K-MARVEL for Short-read sequencing datasets with optimized kmer and extension size (6,9).

Next, we investigated the false positives and negatives for both short and long read datasets. We found that the false positives were mainly categorized into three groups.

The first group comprises ARGs K-MARVEL assigned to a different variant within the correct gene family. This misassignment stems from a fundamental difference in scoring algorithms. Assembly-based approaches rely on protein BLAST, which uses compositional matrix adjustment to calculate alignment bit-score^19^. In contrast, K-MARVEL employs a global alignment to identify the closest reference gene. Because of these two different approaches, the assignment approach makes them different even if the assembled gene is the same in both cases.

The second group comprised ARGs missed by the assembly-based approach but detected by K-MARVEL. These were cases where the assembled gene failed to meet the identity or coverage thresholds pre-defined for the comparison, yet K-MARVEL successfully identified them from the raw sequencing data.

The last category had the least frequency of false positives, emphasizing that these are rare events. Genes falling within this category were fragmented by large insertions in the assembly. Because of this, these genes were filtered out by the assembly-based approaches due to failing the coverage-based threshold. However, these fragmented genes were fully captured by our tool due to the availability of k-mers for the entire gene body. While this is a key strength of K-MARVEL, such detections were classified as false positives because the ground truth data comprised ARGs captured by the assembly-based methods.

Regarding false negatives, most were either mis-assigned by K-MARVEL with respect to the ground truth, or because the sequencing data did not sufficiently cover the target gene.

Detailed information about the false positives and negatives categorized based on sequencing technology is available in the Supplementary Table 4.

### Computational Performance and Scalability

To evaluate the computational efficiency of K-MARVEL, we collected the run-time metrics of the tool across all tested empirical datasets with the optimized k-mer and extension size. The short-read datasets ranged from 1 million to 32 million read pairs, and the long-read datasets ranged from 4,000 to 1.5 million reads. All runs were conducted on a server equipped with AMD EPYC 7543 (32 cores) and 300 GB of RAM, with the tool configured to utilize 16 threads. Execution time was measured as the total wall-clock time (in seconds), and peak memory usage was recorded in GB. For short-read datasets, the execution time ranged from 25.53 seconds to 157.25 seconds, whereas the peak memory usage varied from 0.54 GB to 2.79 GB. In the case of long-read datasets, the execution time ranged from 4.11 seconds to 771.57 seconds, whereas the peak memory usage varied from 0.88 GB to 8.56 GB.

While comparing the execution time with respect to the dataset size (number of reads), we observed two distinct, linear performance profiles corresponding to the sequencing technology used (Supplementary Figure 1). For short-read data, execution time showed a strong and significant correlation with the number of reads in the dataset (Pearson’s R = 0.67, p<0.001). This strong correlation suggests a highly efficient and low computational cost for short-read sequencing data. In contrast, long-read sequencing datasets exhibited a relatively steeper linear relationship (Pearson’s R = 0.62, p<0.001). While the correlation was statistically significant, we observed greater variance in run-time for long-read datasets, which is likely attributed to the read length agnostic technology. We also assess the correlation between computational metrics and data size in terms of the total number of sequenced bases. The tool’s execution time scaled linearly with respect to the number of sequenced bases for both long-read (Pearson’s R = 0.92, *p*<0.001) and short-read (Pearson’s R = 0.63, *p*<0.001) datasets. This near-linear relationship indicates that the run-time grows in a scalable manner (Figure 3A). Regarding memory footprint, the relation strongly correlated with long-read sequencing datasets (Pearson’s R = 0.37, *p*<0.001). However, it showed no significant correlation for short-read datasets (Pearson’s R = 0.007, *p*=0.96). This demonstrates that the tool’s memory footprint remains low and stable regardless of the dataset’s size in the case of short-read sequencing datasets (Figure 3B).

**Figure 3:**
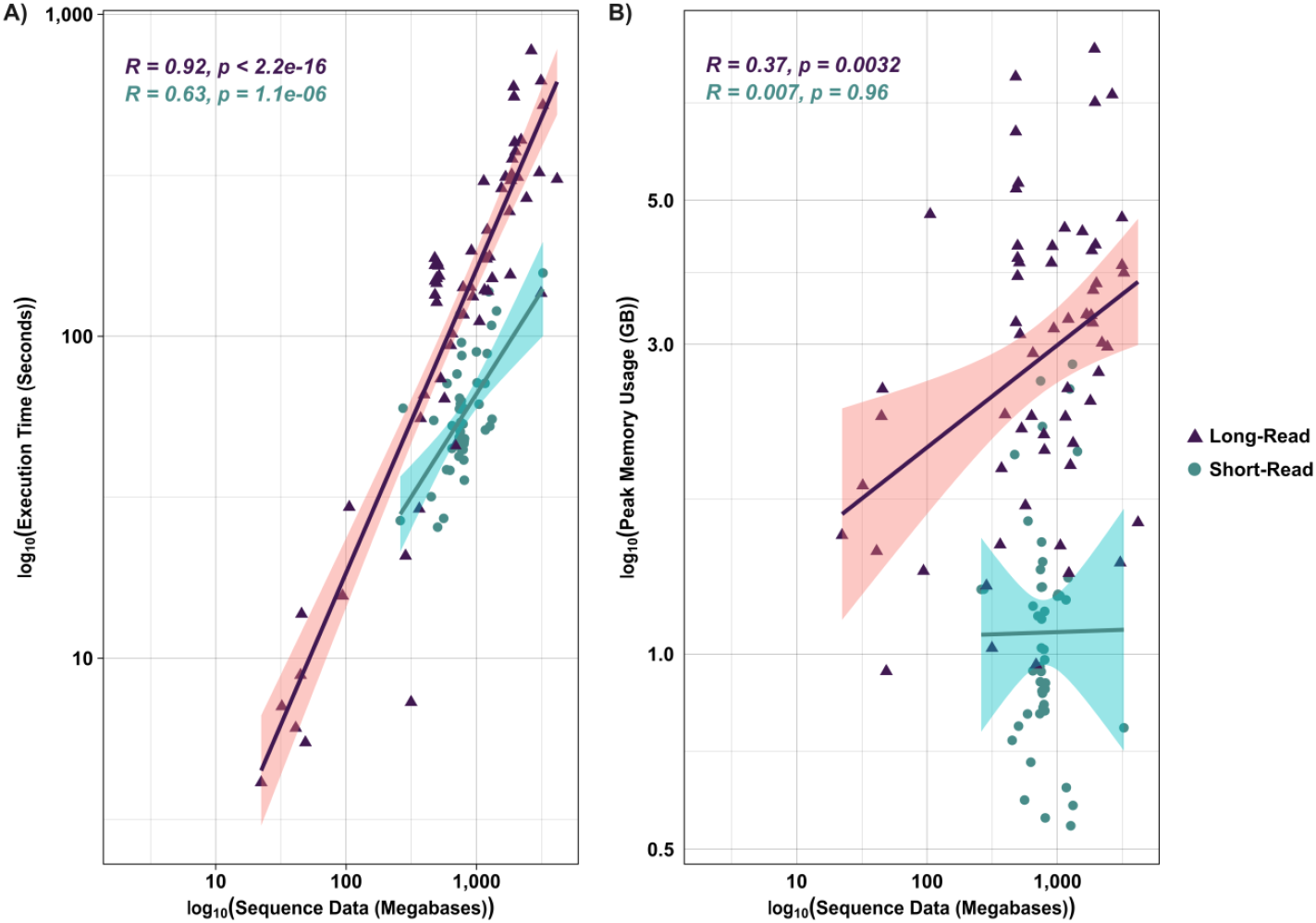
Computational Performance and Scalability of K-MARVEL. The tool’s run-time and peak memory usage were recorded across diverse sets of short-read (teal circles) and long-read (purple triangles) datasets. Both axes are scaled to log10 to visualize performance across several orders of magnitude. Pearson’s R was calculated to assess the relationship between the computational metric and the sequencing data. A) Execution time. B) Peak memory usage. The execution time scaled near-linearly for short and long-read datasets, showing the tool’s scalability. Memory consumption for short-read datasets remains consistently low and stable regardless of the data size; however, it scales linearly for long-read datasets.

Detailed information about the execution time, peak memory usage and other computation metrics is provided in the Supplementary Table 5.

Next, we compared the tool’s computational performance to the most widely used assembly-based approaches. We utilized Flye version 2.9.6^20^ for assembling the long-reads and SPAdes version 4.2.0^21^ with the ‘--isolate’ parameter for assembling the short-reads. RGI version 6.0.3 was then used to capture the ARGs from the assembled contigs. All the tools were tested with 16 computation threads on a server equipped with AMD EPYC 7543 (32 cores) and 300 GB of RAM. K-MARVEL performed significantly faster than the assembly-based approaches for both short and long-read sequencing datasets (Figure 4A), with a median execution time of 151.18 seconds for the long-read as compared to 808.13 seconds for Assembly based approach. For short-read datasets, K-MARVEL achieved a median execution time of 53.68 seconds, which was significantly lower than the Assembly approach (427.4 seconds) (*p-value* < 0.0005, Wilcoxon rank-sum test). The tool’s memory footprint was also significantly lower than the assembly approach for short-read sequencing datasets (Figure 4B). While processing short-read datasets, it maintained a median peak memory footprint of 1.021 GB which was approximately 10 times lower than the memory footprint of the Assembly based approach (10.856) (*p-value* < 0.0005, Wilcoxon rank-sum test). In case of long-read datasets, the comparison was non-significant as K-MARVEL utilized a median of 3.12 GB which was slightly higher than the Assembly based approach (3.02 GB) (*p-value* = 0.38, Wilcoxon rank-sum test), it however, exhibited a tighter distribution with fewer high-memory outliers.

**Figure 4:**
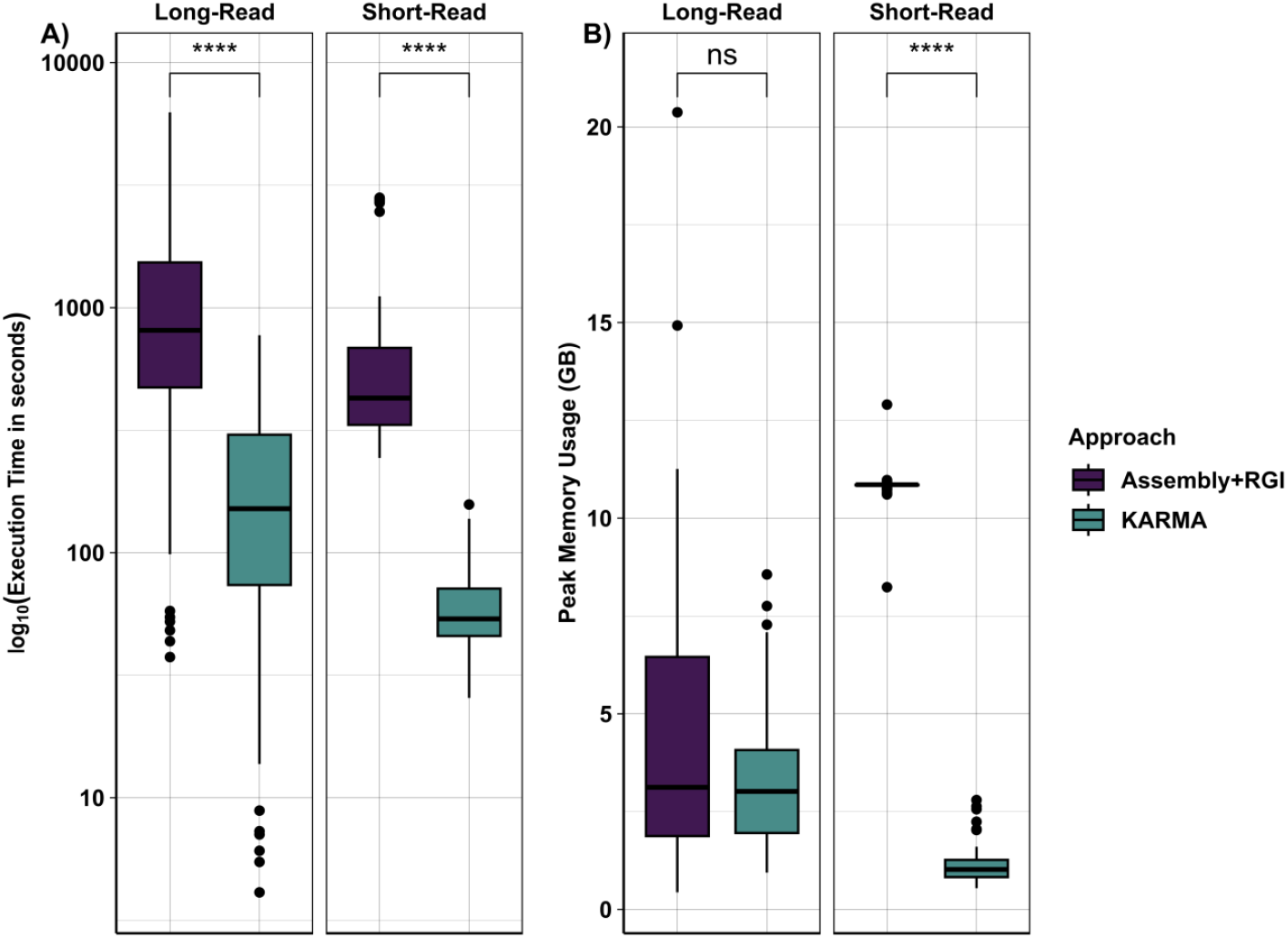
Comparison of run-time performance metrics between K-MARVEL and assembly-based approaches. **A)** Boxplot comparing execution time (seconds) between the assembly-based approach and K-MARVEL. ‘ ****’ indicates p-value ≤ 0.0005; p-value was calculated using the Wilcoxon ranked-sum test. **B)** Boxplot showing the comparison of peak memory usage (GB) between the assembly-based approach and K-MARVEL. ‘ns’ indicates p-value > 0.05 (non-significant) and ‘****’ indicates p-value ≤ 0.0005; p-value was calculated using the Wilcoxon ranked-sum test. K-MARVEL significantly reduces execution time and memory consumption for short-read datasets and execution time for long-read datasets.

Regarding execution time, K-MARVEL performed faster than the assembly-based approaches for both short and long-read datasets. The tool’s memory footprint was significantly lower than its counterpart for short-read datasets. However, it had a comparable memory footprint with the assembly-based approach for long-read sequencing data. Based on these observations, it is safe to say that K-MARVEL is suitable for both long and short-read datasets regarding classification accuracy and computational footprint, making it suitable to process large metagenomic datasets.

## Discussion

In this study, we developed and validated K-MARVEL, a high-performance computational method for capturing the resistant genes and resistance-conferring mutations from raw sequencing data. INDELs errors are particularly common in Oxford nanopore sequencing technology due to homopolymer regions and modified bases in the surrounding k-mers^9,22^. This can lead to false detection of resistance-conferring variants, as it is difficult to distinguish between sequencing errors and true mutations. K-MARVEL overcomes this challenge by doing all its operations in protein space, significantly reducing its execution time and memory footprint. This makes it particularly well-suited for rapid analysis in settings where long-read sequencing is employed for real-time surveillance.

In our testing, K-MARVEL demonstrated superior accuracy and computational efficiency compared to existing methods for both short and long read sequencing technologies. It achieved a high F1-score of 0.9783 and 0.9754 for short and long read sequencing, respectively, indicating a robust balance of Precision and Recall.

The recent advancement of miniature, portable, point-of-care sequencing devices requires efficient, lightweight analysis tools. With the computational performance of K-MARVEL in the case of both short and long read sequencing datasets, it is safe to assume that the tool can be executed efficiently in modern portable laptop computers, making it perfect for portable point-of-care devices.

A further advantage of K-MARVEL is its capacity to reconstruct the full-length resistance genes often fragmented in *de novo* assemblies due to large insertions or repetitive regions. While such fragments may be discarded by assembly-based ARG detection approaches, especially due to the lower coverage, K-MARVEL’s k-mer-based method can reassemble the complete gene by leveraging k-mers that span the entire gene body. For instance, in the *Shigella flexneri* AUSMDU00009397 reference genome assembly, the sul2 gene is split into two fragments (73.06% and 26.94% of the gene). Because of the low coverage, both the fragmented regions were discarded in the assembly-based approaches; however, our tool reconstructed the full gene. This ability to infer complete functional genes from fragmented data provides a more biologically and clinically relevant view of the resistome than an analysis limited by assembled genomic structure.

A recent innovation in nanopore sequencing is duplex base calling, a process that uses template and complement strands to generate a consensus sequence, thereby significantly reducing error rates^23,24^. Although duplex reads constitute a small fraction of total sequencing output compared to simplex reads^25^, our analysis demonstrates that a mere 5,000 duplex reads (∼4x sequencing depth) are sufficient for K-MARVEL to accurately detect antimicrobial resistance genes and resistance-conferring mutations. This finding is critically important for analyzing complex clinical samples, where high levels of host DNA contamination often result in a very low yield of microbial reads. Applying K-MARVEL to duplex basecalled data in such scenarios provides an effective strategy for profiling the resistome.

Despite its advantages, K-MARVEL has some limitations. Because of its core principle of reference k-mer match and extend, it is heavily reliant on the reference gene database. Hence, it is not suited for discovering entirely novel resistance determinants.

Although several well-maintained ARG databases are available, such as MEGARes^26^, the national database of antibiotic-resistant organisms (NDARO)^27^, and ResFinder^28^, K-MARVEL’s current implementation is compatible solely with the comprehensive antibiotic resistance database (CARD)^29^. However, we are actively developing extensions to incorporate additional well-maintained databases.

Finally, while K-MARVEL can reconstruct the complete gene from the fragmented genomic context, it does not characterize the underlying structural variants responsible for the fragmentation; this analysis still requires whole-genome assembly.

## Conclusion

In conclusion, K-MARVEL is a fast, flexible, scalable ARG classifier that can capture both ARGs and resistant chromosomal genes from raw sequencing data. The support for long and short-read sequencing data makes it a potential computational method for translational applications.

## Availability and Requirements

K-MARVEL is implemented in Rust, open-source and freely available at git clone https://bitbucket.org/amr-avenger/k-marvel/src/main/.

No new data has been generated during this study. Details about the publicly available datasets used in the study are included in this published article [and its supplementary information files].

