## Supplementary Figures for "K-MARVEL: K-Mer based Antimicrobial Resistance Virtual Exploration Lab"

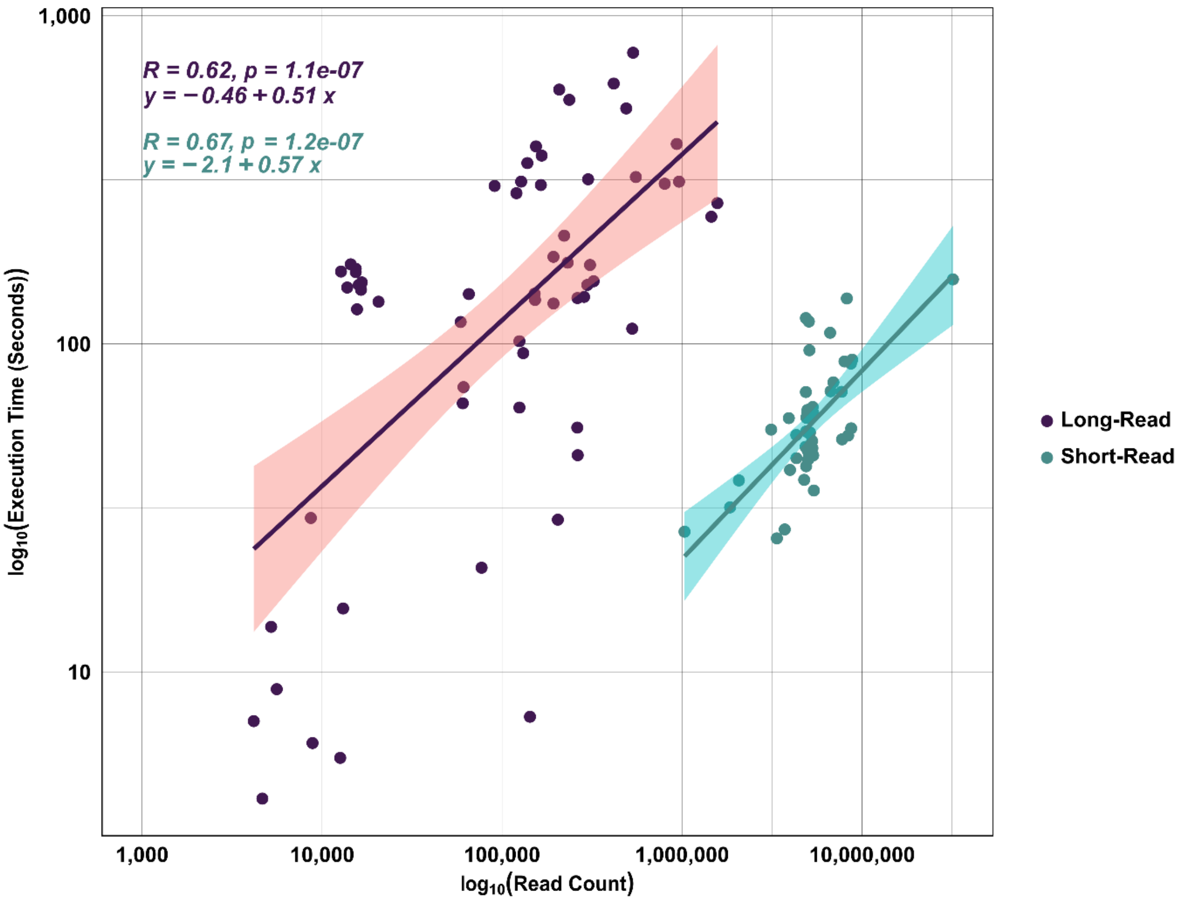


**Supplementary Figure 1:** Computational performance of K-MARVEL. The run-time of the tool was recorded across diverse sets of short-read (teal circles) and long-read (purple triangles) datasets. Both axes are scaled to $\log_{10}$ to visualize performance across several orders of magnitude of sequence data. Pearson's R was calculated assess the relationship between the execution time and the number of reads. The execution time scaled near-linearly for both short and long-read datasets, showing the tool's scalability. While the correlation was statistically significant, we observed greater variance in runtime for long-read datasets, which is likely attributed to the read length agnostic technology.
